# Multi-dimensional transcriptome analysis reveals modulation of cholesterol metabolism as highly integrated response to brain injury

**DOI:** 10.1101/2020.12.29.424680

**Authors:** Victor Gourain, Olivier Armant, Luisa Lübke, Nicolas Diotel, Sepand Rastegar, Uwe Strähle

**Author notes:** Correspondence: Victor Gourain, Uwe Strähle.

## Abstract

**Background:** Zebrafish is an attractive model to investigate regeneration of the nervous system. Despite major progress in our understanding of the underlying processes, the transcriptomic changes are largely unknown. We analysed the transcriptome of the regenerating telencephalon for changes in the expression of mRNAs, their splice variants and investigated the putative role of regulatory RNAs in the modulation of these transcriptional changes.

**Results:** Profound changes in the expression of genes and their splice variants engaged in many distinct processes were observed. As exemplified by the coordinated regulation of the cholesterol synthesizing enzymes and transporters, the genome responded to injury of the telencephalon in a multi-tiered manner with distinct and interwoven changes in expression of enzymes, transporters and their regulatory molecules. This coordinated genomic response involved a decrease of the mRNA of the key transcription factor SREBF2, induction of microRNAs *(miR- 182, miR-155, miR-146, miR-31)* targeting cholesterol genes, shifts in abundance of splice variants as well as regulation of long non-coding RNAs.

**Conclusions:** Differential transcription and splicing are important processes in response to injury of the telencephalon. Cholesterol metabolism is switched from synthesis to relocation of cholesterol. Based on our *in silico* analysis, this switch involves complementary and synergistic inputs by different regulatory principles. Our studies suggest that adaptation of cholesterol metabolism appears to be a key process involved in regeneration of the injured zebrafish brain.

## Introduction

The capacity of the human brain to regenerate damaged tissue is very limited. In contrast, the central nervous system (CNS) of some fish species has a remarkable ability to regenerate with full restoration of functions [1, 2, 3, 92, 93]. For example, deep wounds inflicted by stabbing the brain of the zebrafish with a needle will fully heal. For this [1, 3] and all its other experimental advantages [72, 73], the zebrafish has become a powerful model for the analysis of regeneration of the CNS of vertebrates. Although many brain regions show strong cell proliferation and regenerative responses to injury [44], the telencephlon has emerged as a favoured tissue to study regenerative neurogenesis in zebrafish. The everted structure of the teleost telencephalon [4] presents proliferative cell bodies immediately below the skull and the thin telachoroidea at the dorsal surface [5, 18, 8]. The telencephalic hemispheres are thus easily accessible to wounding and observation without damaging other brain tissues [1 43, 94].

Radial glia cells (RGCs) are the stem cells driving the regenerative response in the zebrafish telencephalon [6, 8, 18]. Their cell bodies are scattered at the periventricular surface from the everted dorsal aspects to the medial areas of the telencephalon. Their thin processes span the entire parenchyma from the cell bodies at the periventricular area to the pial surface. RGCs differentiate into either neuronal cells or, in self-renewing cycles, into more radial glial cells [81]. In response to injury, RGCs undergo mostly symmetric divisions [9, 11]. This type of division is rarely observed in the intact telencephalon during constitutive neurogenesis [15] and may eventually result in a depletion of the neural stem cell (NSC) pool.

Cell death was an immediate reaction to damage of the tissue by 4 hours post lesion (hpl) followed by recruitment of microglia and peripheral immune cells to the lesion [9, 10, 11, 12]. Edema developed at 24 hpl [9]. Oligodendrocytes and oligodendrocyte precursors accumulate at the site of the injury [1] similar to what is observed in the mouse brain. However, in contrast to the mouse brain, oligodendrocytes or their precursor did not proliferate significantly in the zebrafish [1, 11]. By 48 hpl, the RGCs start to divide at a higher rate above the baseline levels characteristic of constitutive neurogenesis [1, 18, 43, 98]. When only one hemisphere of the telencephalon is injured, this proliferative response as well as gene activation is entirely restricted to the injured hemisphere [1, 14]. Thus, signals increasing proliferative responses are limited to the injured hemisphere despite the close juxtaposition of the ventricular surfaces in the medial region of the telencephalon. Proliferation of RGCs reaches a peak at 6 to 8 days after lesion (dpl) and then decreases steadily again reaching basal levels by 10 days [1, 13, 98]. This proliferation of RGCs can be triggered by inflammatory signals [12].

Transcription is a tightly regulated process, where cross-talk between epigenetic marks, transcription factors and their *cis*-regulatory elements orchestrate gene expression. On top of these complex interconnected *cis-* and *trans*-regulatory processes, alternative splicing offers an additional layer to modulate transcriptional responses by increasing the functional diversity of proteins by exon inclusion or exclusion or affecting the stability of mRNAs and proteins [28]. Expression levels are further fine-tuned by regulatory RNAs (microRNAs (miRNAs) and long non-coding RNAs (lncRNAs)). Measuring changes in the repertoire of spliced isoforms and key regulators in relation to differentially expressed gene ontology groups can help deciphering the molecular processes underlying brain regeneration.

Previously, we identified by deep sequencing 252 transcription factor (TF) genes which were up-regulated and 27 TF genes that were down-regulated upon injury [14]. The expression pattern of these genes was mapped together with 1,202 constitutively expressed regulators of transcription [13, 14]. These previous studies focused on the response of transcription factor genes to injury and repair of the telencephalon. Here, we have broadened the analysis of our RNASeq data to all gene ontologies to identify pathways and biological processes that are activated or repressed in response to injury. Besides the expected processes such as neurogenesis and axonal growth, we identified, among many others, genes related to cholesterol metabolism to be differentially expressed in response to injury. This response was multi-tiered and highly coordinated. While mRNAs encoding synthesizing enzymes were down-regulated, transporters were up-regulated. Moreover, transcriptional changes indicated regulation of expression at multiple levels, from the down-regulation of the master TF of cholesterol synthesising enzymes, Srebf2, to the up-regulation of miRNAs with target sequences in cholesterol synthesizing enzymes and Srebf2 itself. Finally, mRNAs of cholesterol transporters and synthesizing enzymes were differentially spliced suggesting alternative splicing as yet another mechanism for fine-tuning cholesterol metabolism. Our data strongly suggest that modulation of cholesterol metabolism is a key process in brain regeneration in the zebrafish. In addition, our study provides the first comprehensive analyses of basal and injury induced expression of miRNAs and long non-coding RNAs and the shifts in splice patterns in the transcriptome of the regenerating zebrafish telencephalon. We thus report here also valuable resources for follow-up studies.

## Results

### Injury-induced changes in steady state levels of polyadenylated RNAs in the telencephalon

To get a comprehensive picture of the transcriptional changes caused by injury of the adult brain, we re-analysed the previously established RNASeq data [14]. The sequenced cDNA was derived from polyadenylated RNA isolated from injured telencephala of the adult zebrafish at 5 dpl, with the contralateral hemisphere as uninjured control [14]. We analysed in total approximately 600,000,000 reads from injured telencephalic hemispheres and an equal number of reads from uninjured control hemispheres. The RNAseq samples from the three biological replicates of each condition were consistent as assessed by hierarchical clustering (Figure 1A). A total of 32,520 genes annotated in the zebrafish reference genomes GRCz11 were tested and 17,301 were expressed in the adult zebrafish telencephalon (Figure 1B). The analysis of differential expression revealed 1,946 and 3,043 genes with significantly up- or down- regulated expression respectively (adjusted p-value (adjp)< 0.05), (Figure 1B, Supplementary Table 1) relative to the transcriptome of the uninjured hemisphere.

**Figure 1:**
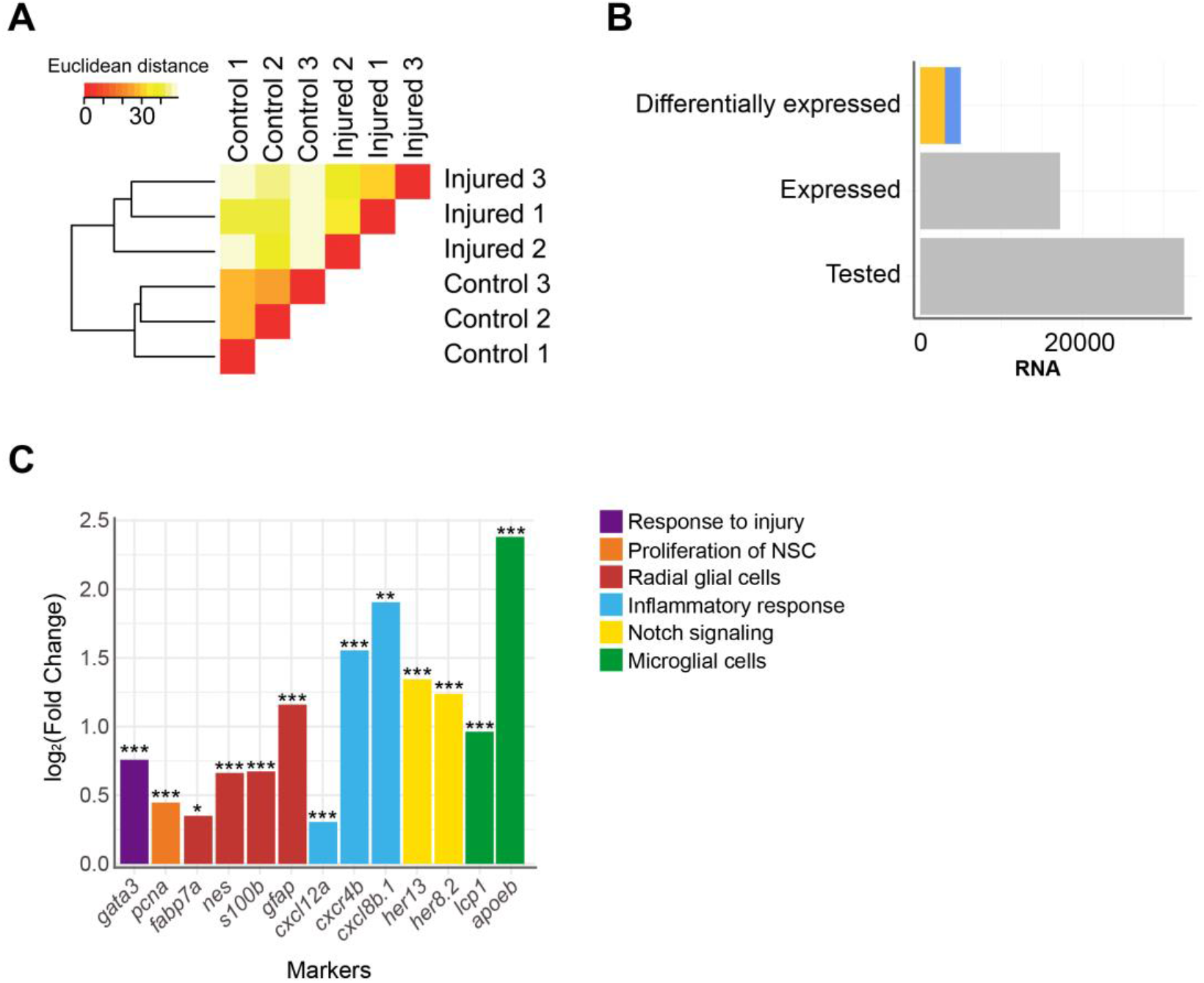
Injury-induced changes in level of polyadenylated RNAs. **A** Hierarchical clustering of RNASeq samples. The heat map represents the Euclidean distance between samples. **B** Quantification of RNAs analyzed by RNASeq. yellow: increased level after injury and blue: decreased level after injury. **C** Markers with significant changes in level of transcript. * adjp<=0.05, ** adjp<10^-02^, *** adjp<10^-04^. Color code indicates different gene ontology terms.

To assess the sensitivity of our analysis, we selected genes known from previous studies to be altered in their level of expression by injury of the telencephalon (Figure 1C). The transcription factor *gata3* is a gene which responds to injury of the telencephalon very rapidly [43]. In agreement, the level of transcripts coding for *gata3* was significantly increased upon injury (Fold Change (FC)=1.70; adjp<10^-03^). Similarly, transcripts coding for *proliferation cell nuclear antigen* (PCNA), a marker of dividing cells [17] were elevated after injury (FC=1.37; adjp<10^-04^), as well as mRNAs of the RGC-specific genes *fabp7a, nestin, s100b, glial fibrillary acidic protein (gfap)* [18, 7] (FC=1.27, 1.58, 1.59 and 2.23, respectively; adjp<0.05, <10^-05^, <10-^05^ and <10^-24^, respectively). We also observed mRNAs encoding Apoeb and Lcp1, markers for microglia [19], were up- regulated upon injury (FC=5.21 and 1.95, respectively; adjp<10^-67^ and <10^-06^, respectively) as were mRNAs of the cytokines *cxcl8b.1* and *cxcl12a* (FC= 2.93 and 1.23, respectively; adjp<10^-35^ and <10-^03^, respectively) and the cytokine receptor *cxcr4b* (FC=3.73; adjp<10^-02^). The increased expression of these genes coding for cytokines and cytokine receptors reflects the activation of an inflammatory response by injury [12]. Changes in genes promoting quiescence of progenitors were also detected, for instance the level of transcripts coding for the Notch target TFs Her13 and Her8.2 [97] were increased after injury (FC=2.53 and 2.36, respectively; adjp<10^-21^ and <10^-06^, respectively). Taken together, all assessed genes whose expression levels are known to be regulated by injury were verified in our transcriptome analysis (Figure 1C). These results show that we detected variation of transcripts levels in response to telencephalon injury with high sensitivity.

### Gene ontology analysis

Next, we assessed the enrichment of specific ontologies among regulated genes to obtain information on the biological processes that are linked to the repair of the injured telencephalon. Three sources of data provided information on biological functions (Gene Ontology (GO), [20]), signalling pathways (Kyoto Encyclopedia of Genes and Genomes (KEGG), [21)] and metabolic pathways (Reactome, [22]). A total of 192 GO terms, 34 KEGG pathways and 295 Reactome enzymatic reactions were significantly enriched among the genes with variation in level of transcripts upon injury (adjp<0.05)(Supplementary Table 2).

A major response was “generic transcription regulation” comprising, among others, the GO terms “RNA polymerase II transcription factor activity” (associated with 63 differentially expressed genes; adjp<10^-02^) and “Regulation of transcription” (represented by 230 differentially expressed genes; adjp<10^-21^). This broad response of the transcription regulator genes is consistent with previous reports [13, 14] and reflects the large-scale, injury inflicted changes of the transcriptome. Other GO terms - expected from the response of the genome - were “neurogenesis”, “angiogenesis”, “immune response” (Figure 2). Beside these expected functions among the regulated genes, we detected a large number of enriched gene ontology terms such as “mRNA splice site selection” (adjp=0.025) and “cholesterol metabolism” (adjp<10-^04^).

**Figure 2:**
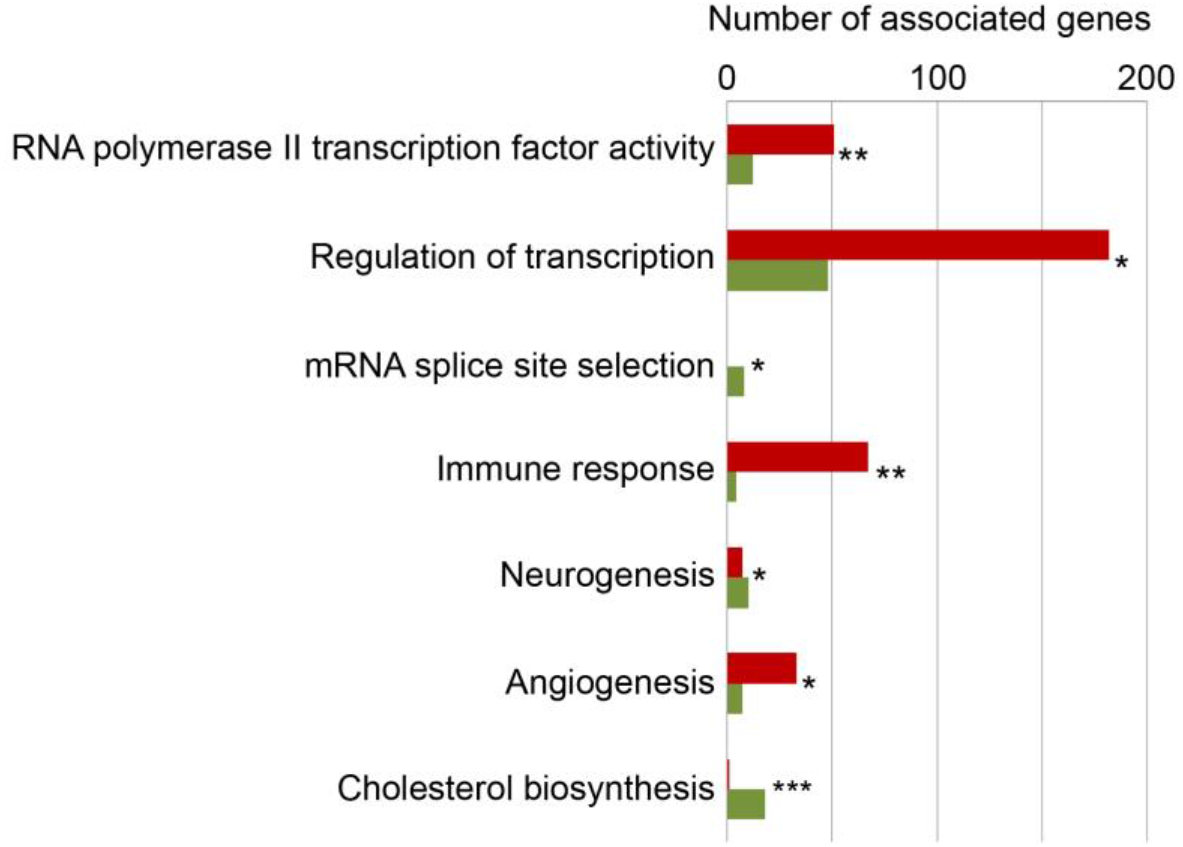
Selected enriched biological functions. * adjp<=0.05, ** adjp<0.01, *** adjp<0.001; red: up-regulated genes; green: down-regulated genes. (For complete list see Supplementary Table 2)

### Expression of enzymes and transporters of cholesterol metabolism are coordinately regulated in response to injury

Since nothing is known about a role of cholesterol in the regeneration of the zebrafish brain, we focused our analysis on genes linked to cholesterol metabolism. Cholesterol is a component of plasma membranes and is highly abundant in myelin sheaths [50]. The synthesis of cholesterol from acetyl-CoA is a multi-step process involving a large number of distinct enzymes [26]. We found that, after injury, levels of transcripts coding for all but one of the enzymes involved in cholesterol synthesis were decreased (Figure 3A and Supplementary Table 3). Cholesterol synthesis is repressed by availability of external cholesterol [24]. Thus, besides synthesis, transport of cholesterol is an important process that contributes to the regulation of cholesterol levels. Upon brain injury, levels of mRNAs coding for 8 cholesterol transporters were significantly increased (Figure 3A; Supplementary Table 3). Taken together, our results indicate that cholesterol transport is increased while its synthesis is turned down. We also observed that the level of transcripts of the growth factor *brain-derived neurotrophic factor* (*bdnf*) known to repress cholesterol synthesis and to promote together with ApoE and ABCA1 cholesterol transport [51] was significantly decreased upon injury (FC=0.87; adjp=0.031). Together, these data suggest that cholesterol transport and cholesterol synthesis are regulated in a coordinated way in the injured telencephalon.

**Figure 3:**
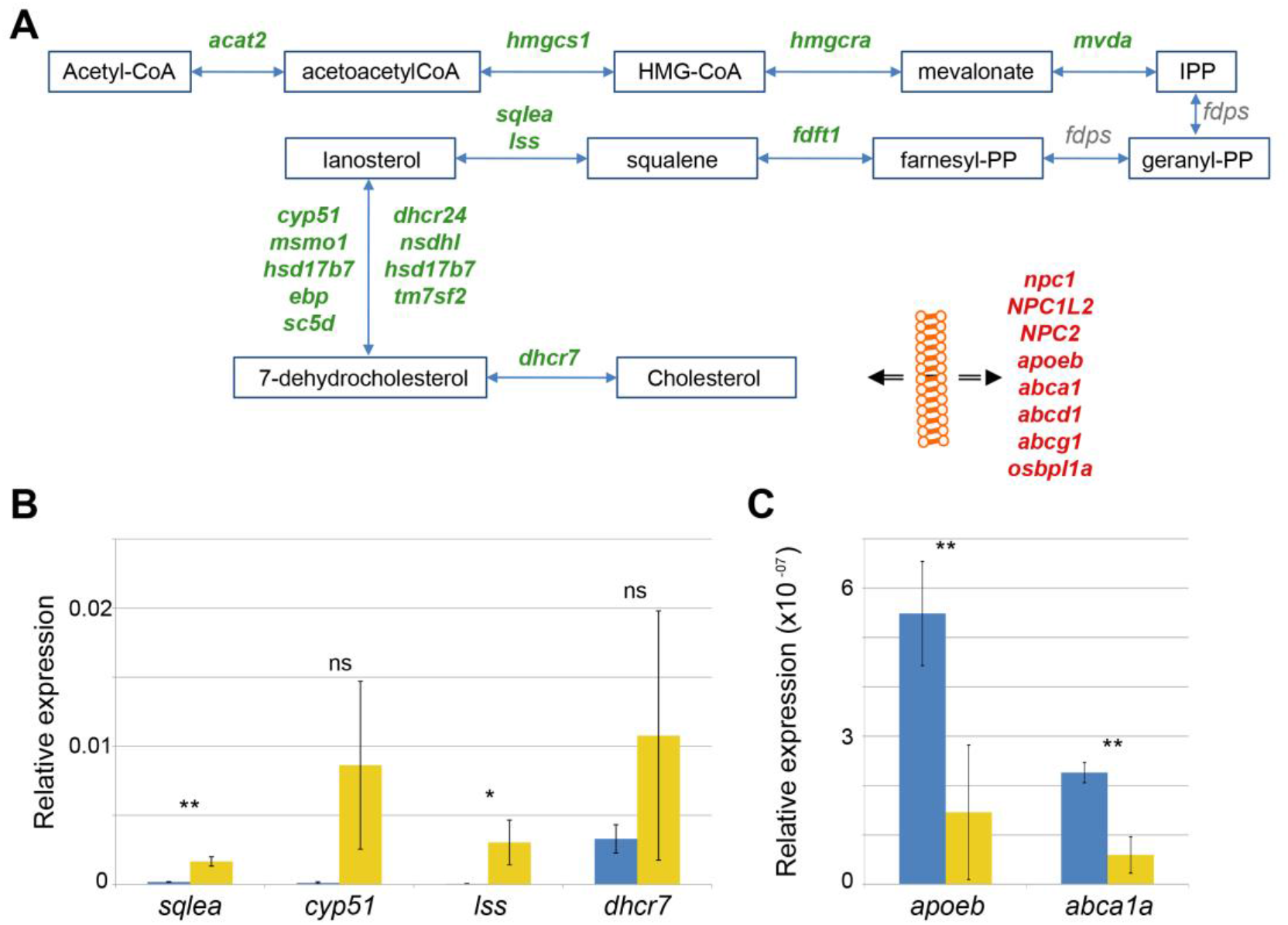
Alteration in cholesterol metabolism in response to brain injury. **A** Cholesterol synthesis pathway. red: up-regulated genes, green: down- regulated genes. Products and substrates are represented in blue boxes and enzymatic reactions by blue arrows. Bottom right: transporter proteins involved in ferrying cholesterol through the body and across membranes. While cholesterol synthesizing enzymes are down-regulated, transporters are up- regulated in the regenerating telencephalon. **B-C** Quantification of mRNAs encoding enzymes synthesizing cholesterol (**B**) and transporters (**C**) using qRT- PCR. * p-value<=0.05, ** p-value<10^-03^, yellow: injured telencephalic hemisphere, blue: control telencephalic hemisphere.

Although our RNASeq transcriptome analysis detects genes responsive to injury with a high sensitivity and fidelity, we wished to confirm these findings on the cholesterol pathway with an independent method and with different sample preparations. To this end, we carried out a qRT-PCR analysis with a number of selected genes (Figure 3B, C). The metabolic enzymes Sqlea, Cyp51, Lss and Dhcr7 yielded lower signals relative to the uninjured control (Figure 3B) as expected from the transcriptome analysis. Similarly, we detected increases of transporter cDNAs encoding *apoeb* and *abca1a* in the injured sample relative to uninjured control cDNA (Figure 3C). These qRT-PCR results verify our transcriptome analysis and support the hypothesis that cholesterol metabolism is modulated after telencephalon injury. Taken together, this response of the transcriptome suggests that injury results in an increase of available cholesterol, presumably as a result of release from damaged and dying cells.

### Expression of the master regulator of cholesterol synthesizing enzymes Srebf2 is reduced upon injury

Basic helix loop helix SREBF transcription factors regulate expression of cholesterol synthesizing enzymes in mammals [24, 25]. We thus explored expression of *srebf.* In the zebrafish genome as in that of mammals, two paralogous genes encode the related Srebf1 and Srebf2 genes, and both are expressed in the adult zebrafish telencephalon. The level of transcripts coding for Srebf2 was significantly lower (FC=0.63; adjp<10^-09^) in the transcriptome of the injured zebrafish telencephalon, consistent with the observed lower expression of Srebf2-targeted genes encoding cholesterol synthesizing enzymes. These data suggest that Srebf2 might be the main regulator of cholesterol synthesis in zebrafish as in mammals [26].

We analysed thus next the promoters of the genes differentially expressed after injury for potential enrichment of Srebf2 binding sites. Srebf2 interacts with a short target sequence, the Sterol Regulatory Element (SRE), in the promoter region of responsive genes [26]. We analysed whether the two related consensus sequences of mammalian SRE motifs listed in JASPAR (Figure 4A; left and middle panel) were present in the 1-kilobase (kb) promoter sequence of the genes with significant variations in level of transcripts after injury. As a control, we created a set of background 1-kb sequences obtained from randomly chosen genes.

**Figure 4:**
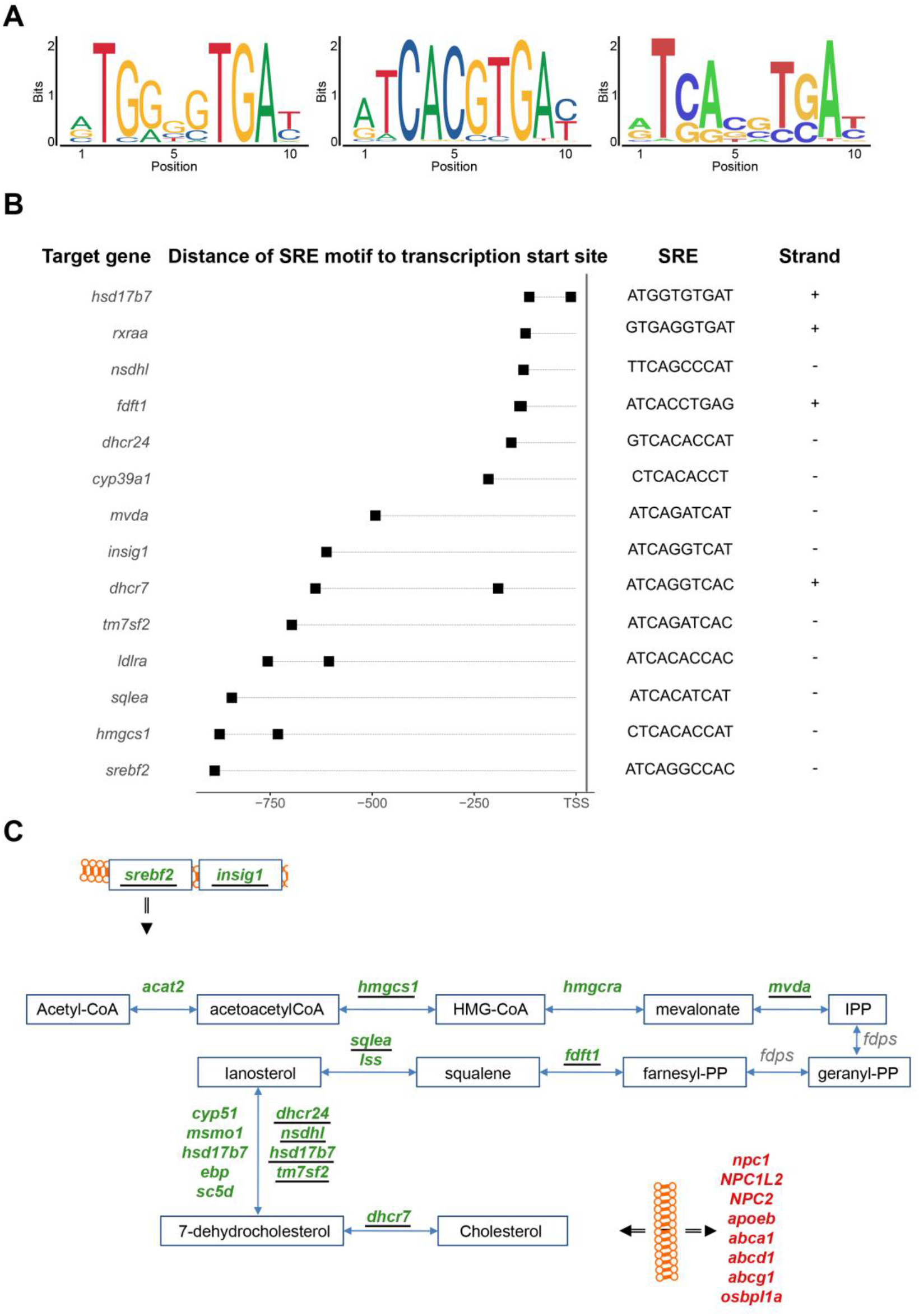
Sterol Regulatory Element (SRE) motif analysis. **A** Human SRE motifs derived from ChIPSeq (left) and Selex (middle) and derived SRE motif from mapping these consensus sequences in the zebrafish genome (right panel). **B** Identification of the SRE motifs in the 1-kb promoter of genes involved in cholesterol metabolism. TSS: transcription start site; +: forward; -: reverse. **C** Genes (underlined) coding for proteins of the cholesterol synthesis pathway harboring a SRE motif in their promoters. Double black arrow: flow across membrane; gene names in green: gene down-regulated; in red, gene up- regulated in response to injury. For further details see also legend to Figure 3A.

In total, 1,145 genes with changes in expression levels upon injury harbour homologies of a SRE motif in the 1-kb promoter region. Relative to the control, this represents a significant enrichment with positive log odds scores and after correction for GC content and repeat of k-mers (Supplementary Table 6). Moreover, the GO term “cholesterol biosynthetic process” is enriched among these genes carrying a SRE motif (adjp<0.05) (Supplemental Table 9). By additionally manually mining the list of SRE harbouring genes, in total nine genes coding for enzymes involved in the synthesis of cholesterol: *hmgcs1, mvda, fdft1, sqlea, tm7sf2, nsdhl, dhscr24, hsd17b7* and *dhcr7* (Figure 4B; Supplementary Table 4) were found with SRE motifs in the 1-kb promoter region. These results partially overlap with SRE motifs mapped in the promoters of the human and mouse orthologous genes [26] (Supplementary Table 4). SRE motifs were also identified in the promoter region of two key regulators of the cholesterol metabolism, *srebf2* itself and *insig1,* a post-translational regulator of Srebf2 [45]. (Figure 4C). The presence of a SRE binding site in the promoter of Srebf2 suggests an auto-regulatory feedback-loop of *srebf2*. The SRE motifs were also identified in the promoter of other differentially expressed genes involved in cholesterol metabolism (Figure 4B). For example, low-density lipoprotein receptor *(ldlra),* the alpha sub-unit of the retinoic X acid receptor *(rxraa)* and cytochrome P450 family 39 subfamily A polypeptide 1 *(cyp39a1),* all involved in cholesterol metabolism, were detected as potential Srebf2 transcriptional targets. From the homology scores in the zebrafish genome (Figure 4A; left and middle panel) [36], a putative zebrafish Srebf2 sequence was derived (Figure 4A; right panel). The *in silico* predicted sequence is similar to the SREBF2 binding sequence identified in human genes by Selex [77] rather than the Chromatin Immuno-Precipitation (ChIP) followed by Sequencing [41] derived consensus sequence (Figure 4A; middle panel).

Taken together, this significant enrichment of SRE motifs in cholesterol biosynthethic genes supports the notion that Srebf2 is also a regulator of the expression of these genes in the zebrafish genome.

### miRNAs that target cholesterol genes are increased upon injury

miRNAs are well established negative regulators of coordinated gene programs [53]. The changes in expression of miRNAs were thus investigated by small RNASeq in the injured telencephalon in comparison to the uninjured brain. Computation of Euclidean distances and hierarchical clustering between small RNASeq samples grouped the samples according to their respective experimental condition (Figure 5A). A total of 184 miRNAs annotated in the zebrafish reference genome (GRCz11) were detected in the transcriptome of the adult zebrafish telencephalon. The analysis of differential miRNA expression, identified 31 miRNAs regulated at least two-fold after injury (adjp<0.05). Among these, the level of 22 miRNAs increased upon injury while the level of 9 miRNA decreased (Figure 5B) (Supplementary Table 7). For further analysis, we focused on the five miRNAs with the strongest variation in their level in response to injury. The level of four miRNAs increased in response to injury: *miR-31* (FC=4.92; adjp<10-^64^), *miR-146a* (FC=4.50; adjp<10-^62^), *miR-155* (FC=2.58; adjp<10^-09^) and *miR-182* (FC=2.28; adjp<10-^02^). The level of *miR-26b,* decreased after injury (FC=0.0050; adjp<10^-246^). None of these five miRNAs were previously shown to be involved in the regulation of constitutive or regenerative neurogenesis.

**Figure 5:**
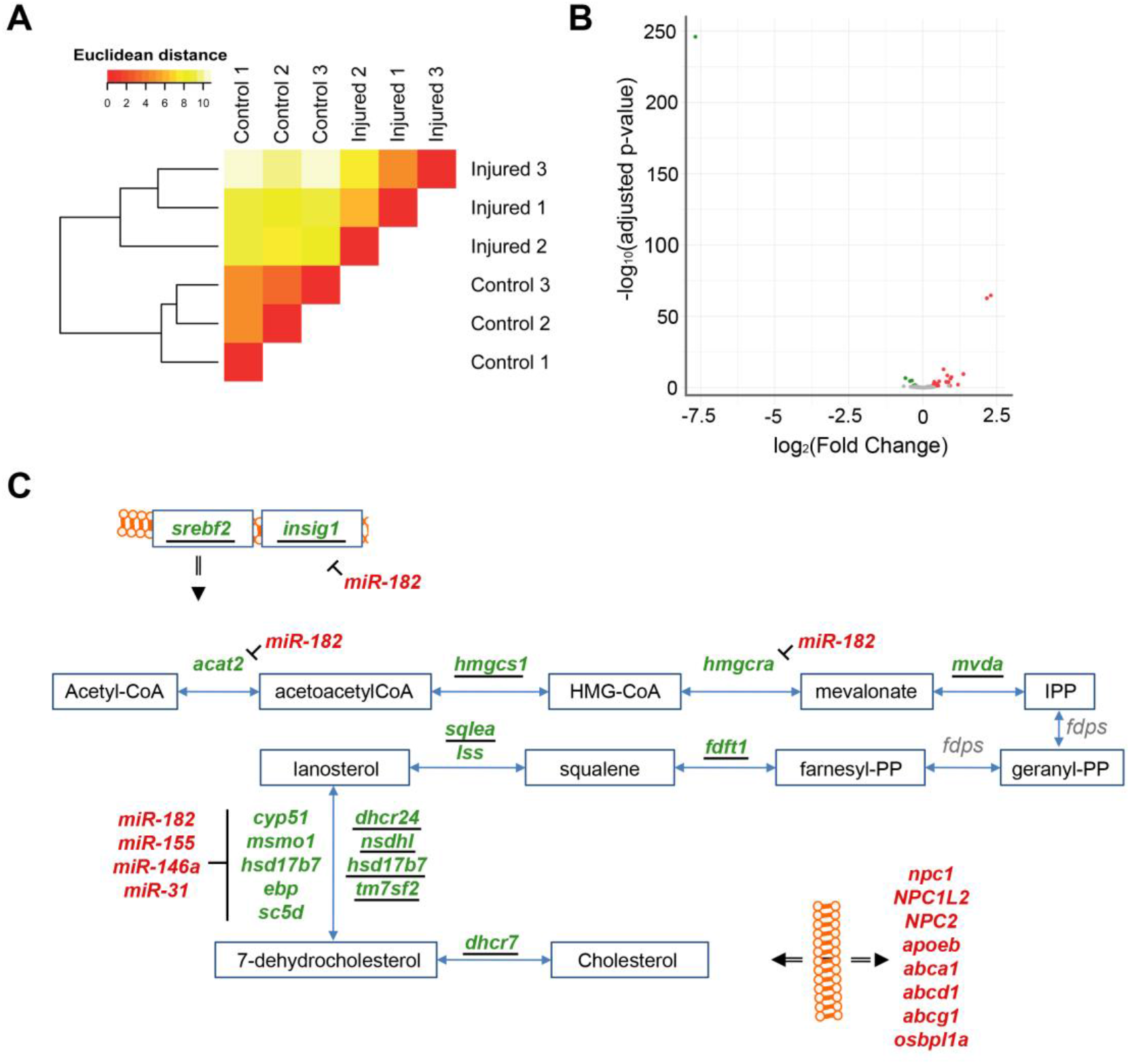
Injury-induced changes in levels of miRNAs. **A.** Hierarchical clustering of small RNASeq samples. **B** Identification of differentially expressed miRNAs. red: up-regulated, green: down-regulated, adjp<0.05. **C** Scheme of cholesterol synthesis with targets of *miR-182, miR155, miR-146a* and *miR-31* indicated. Gene names in green: gene down-regulated; in red gene up-regulated in response to injury. For further details see also legend to Figure 3A and 4C.

We next assessed potential mRNA targets of these five miRNAs by screening for the presence of the seed sequence in the 3’UTR of differentially expressed mRNAs. Interestingly, we found the threemiRNAs*miR-31*, *miR-146a* and *miR-155* target them RNAs of five down-regulated genes coding for enzymes of the synthesis of 7-dehydrocholesterol: *ebp*, *cyp51, sc5d, hsdl7d7* and *msmo1* (Figure 5C). In addition, the mRNAs encoding Insig1 (FC=0.43; adjp<10^-23^), Acat2 (FC=0.75; adjp<10^-06^), Dhcr24 (FC=0.57; adjp<10^-05^), Sc5d (FC=0.66; adjp<10^-03^) and Hmgcra (FC=0.54; adjp<10^-12^) were predicted targets of *miR-182* (Figure 5C). Acat2, Dhcr24, Hmgcra and Sc5d are enzymes participating in the synthesis of cholesterol [26] and Insig1 is a co-factor of Srebf2. Taken together, these data strongly suggest that, in addition to the transcriptional regulation via SREBF2, several miRNAs contribute to the adaptation of the cholesterol metabolism to the altered physiological needs of the injured telencephalon.

### Injury-induced changes in levels of polyadenylated long non-coding RNAs

The vast majority of the known lncRNAs are polyadenylated [16]. Their expression levels can thus be extracted from our RNAseq data. After injury of the adult zebrafish telencephalon, we detected significant changes in the levels of 149 lncRNAs (77 increased and 72 decreased) (Supplementary Table1). As the functional annotation of lncRNAs is still poor, we scored the putative target protein-coding genes next to the loci encoding lncRNAs, and carried out functional annotation enrichment on these nearby protein-coding genes.

Several lncRNAs with changed levels in the regenerating telencephalon were identified directly upstream or downstream of cholesterol-related protein-coding genes (Figure 6). The level of both *oxr1a* lncRNAs and its potential downstream target *sqlea*, known to convert squalene to lanosterol during cholesterol synthesis [26], significantly increased upon injury (Figure 3A). Other examples of potential lncRNA transcriptional target include *pck9* and the lncRNA, *dsg2.1* which were down and up-regulated, respectively. Pcsk9 is known to regulate cholesterol homeostasis [71]. Finally, although no significant change in level was observed for mRNAs coding for *scap,* the level of surrounding lncRNA BX511123.2 significantly changed in response to injury (Figure 6). Scap is a chaperone of Sreb transcription factors and forms a retention complex in the membrane of the endoplasmic reticulum [74].

**Figure 6:**
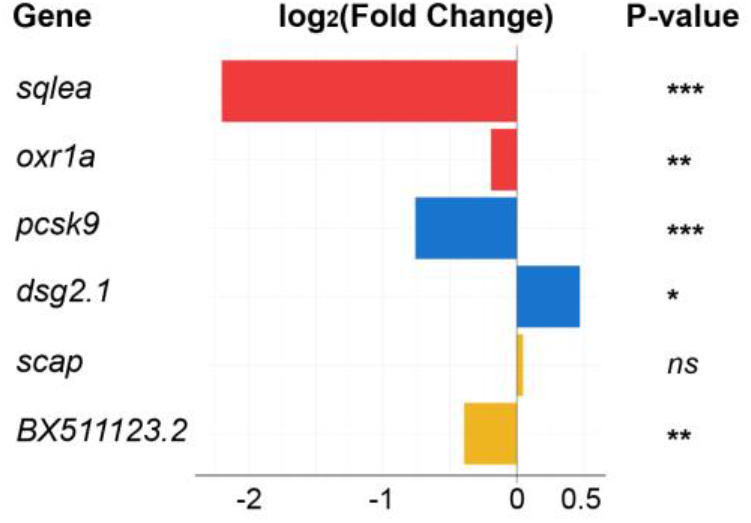
Differentially expressed lncRNAs selected for their association with cholesterol synthesis or transport. **A** Paired non-coding and coding RNAs with changed levels. Color indicates pairs. * adjp<=0.05, ** adjp<10^-02^, *** adjp<10’^04^, ns: not significant. See legend of Figure 7B for the position of the genes in the cholesterol pathway.

Although a regulatory role of any of the lncRNAs has not been established by functional experiments, our data support the hypothesis that lncRNAs are involved in orchestrating the response of the genome to injury of the telencephalon and that they may also more specifically contribute to the regulation of cholesterol metabolism.

### Alternative splicing of RNAs in response to injury of the telencephalon affects cholesterol metabolising enzymes and transporters

Alternative splicing is a post-transcriptional modification of RNAs that increases the functional diversity of proteins by exon inclusion or exclusion or affects the stability of mRNAs and proteins [28]. The enriched gene ontology terms among regulated genes included “mRNA splice site selection” (adjp<0.05) suggesting that injury may alter the pattern of splicing of mRNAs (Supplementary Table 2). Alternative splicing events were identified by comparing *de novo* reconstructed transcripts present in uninjured and injured telencephalic hemispheres taking the annotation of the zebrafish reference genome (GRCz11) into account. FC and adjp were computed for each alternative splicing event. In total, 4,610 alternatively spliced variants were detected in response to injury (adjp<0.05), affecting 1,309 genes. Change of ratio of transcript isoforms was the most recurrent difference between uninjured and injured telencephalic hemispheres. We also identified novel isoforms of RNAs specific for the adult zebrafish telencephalon and which had not yet been annotated in the zebrafish reference genome (GRCz11) (Supplementary Table 8). Thus, brain injury results in a large change of splicing patterns.

These results were further refined according to the biological functions of genes from which alternatively spliced RNAs were synthesized. *Mbpa* (FC=2.6; adjp<10^-08^) and *mpz* mRNAs (FC=7.08; adjp<10^-03^) were alternatively spliced upon injury. These two genes code for components of the myelin sheath [29]. mRNAs encoded by *col12a1a* (FC=4.5; adjp<0.05), *mcamb* (FC=3.59; adjp<0.05) and *myo9aa* (FC=3.6; adjp<0.05) genes were also alternatively spliced after injury. These three protein-coding genes were associated with the development and the regeneration of axons (GO term). Transcripts synthesised from the transcription factor gene *nfia* were alternatively spliced comparing uninjured and injured telencephalic hemispheres (FC=2.24; adjp<0.05). The chicken homologue of *nfia* was implicated in the regulation of gliogenesis in the central nervous system [30]. In response to telencephalon injury, the level of *nfia* transcripts decreased (FC=1.25; adjp<0.05), as well as levels of transcripts of its partners *sox9a* (FC=1.20; adjp<10^-03^) and *sox9b* (FC=1.60; adjp<10^-17^) (Supplementary Table 1).

We next focused specifically on the splicing patterns of genes involved in cholesterol metabolism (see Supplementary Figure for structures of spliced isoforms). In response to injury, the levels of mRNAs encoding the two related zebrafish splicing factors Ptbp1a and Ptbp1b significantly increased (FC=1.57 and 1.20, respectively; adjp<10^-08^ and <0.05, respectively) (Supplementary Table 1). At the same time, the level of *hmgcs1, fads2* and *pcsk9* mRNAs decreased (FC=1.60, 0.66 and 0.59, respectively; adjp<10^-06^, <10^-06^ and <10^-04^, respectively) (Supplementary Table 1). These three genes are all involved in cholesterol metabolism in mammals [46, 47] and *hmgcs1* mRNAs was alternatively spliced in response to injury (adjp<10^-04^) (Figure 7A). A decrease in the number of mapped reads spanning the longest 5’UTR (ENSDARE00001157036, FC=1.41) was consistent with a significant increase in the number of reads spanning the shortest 5’UTR (ENSDARE00001149813, FC=0.18). Drawing from the literature [48], these results suggest that, in response to injury, Ptbp1a/bare involved in the alternative splicing of the 5’UTR of *hmgcs1* transcripts. This likely results in unstable isoforms thus contributing to the reduction of *hmgcs1* mRNA levels in the injured telencephalon.

**Figure 7:**
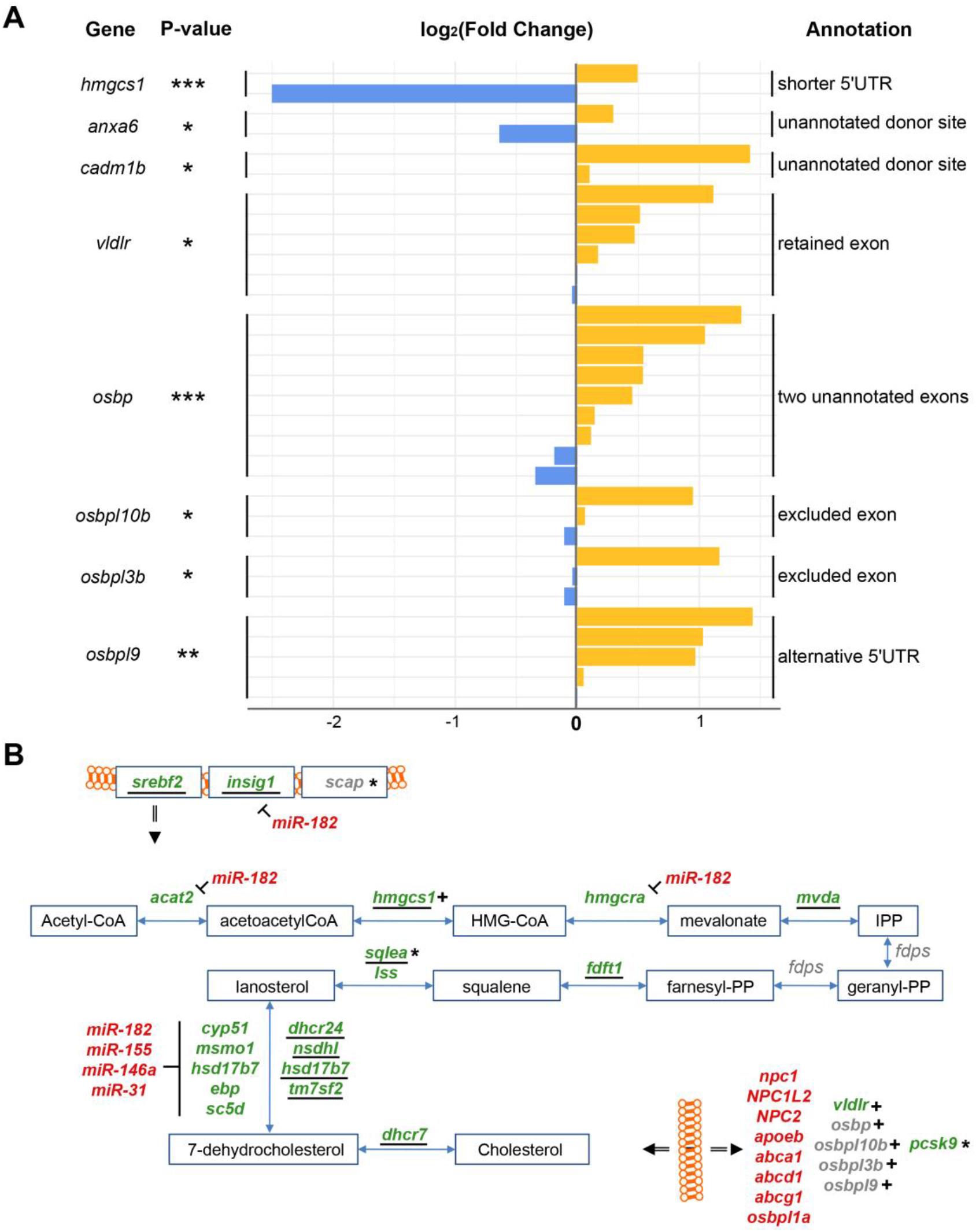
Alternative splicing of RNAs related to cholesterol metabolism in response to injury. **A** Summary of changes in different splice isoforms of the cholesterol-related genes indicated. A number of splice isoforms were not yet annotated in the genome (unannotated). * adjp<=0.05, ** adjp<10^-02^, *** adjp<10^-03^, blue: decrease in number of supporting pairs of reads, yellow: increase in number of supporting pairs of reads. **B** Outline of the enzymes and transporters (bottom right) involved in cholesterol metabolism affected by alternative splicing (indicted by +) and by other modes of possible regulation (miRNAs, lncRNAs). lncRNAs predicted targets (indicated by ^**\***^).Gene names in green: gene down- regulated; red: gene up-regulated in response to injury; underlined: gene with Srebf2 binding site(s) in 1-kb promoter

mRNAs encoding proteins involved in cholesterol transport were also alternatively spliced after injury (Figure 7A). mRNAs encoding the Low Density Lipoprotein Receptor *vldlr* (adjp<0.05) were spliced to exclude an exon (ENSDARE00001166020). No specific protein domain/function was annotated to this exon (InterPro) [55]. LDLs are responsible for extracellular cholesterol transport through the blood stream [56]. Interestingly in contrast to all other cholesterol transporters, the overall level of *vldlr* transcripts significantly decreased upon injury (FC=1.12; adjp<0.05). Two non-annotated splice sites were discovered in exons of *anxa6* (ENSDARE00000906781, FC=0.64 and 1.23, adjp<0.05) and *cadm1b* (ENSDARE00000873208, FC=1.07 and 2.66, adjp<0.05). Anxa6 participates together with NPC proteins in the endosomal trafficking of cholesterol [49], and Cadm1b has a predicted cholesterol 24-hydroxylase activity (GO term).

A total of four mRNAs encoding transporters of cholesterol metabolites of the OxySterol Binding (OSB) family [57] were also affected by splicing in response to telencephalon injury (Figure 7A). Two unannotated exons of *osbp* were discovered as newly emerging upon injury (adjp<10^-05^). In response to injury, an exon was retained in mRNAs encoding *osbpl10b* (ENSDARE00000815047, adjp<0.05) and *osbpl3b* (ENDARE00001041526, adjp<0.05). No corresponding protein domain was annotated (InterPro). Two isoforms of mRNAs encoding Osbpl9b were alternatively spliced in response to injury, including an alternative 5’UTR (ENSDARE00000991106, adjp<0.01) and a retained exon (ENSDARE00001127062, adjp<0.01).

Taken together, our analysis identified alternative splicing as an important response to damage of the zebrafish telencephalon, suggesting distinct isoforms of proteins involved in the repair of the damaged tissue. The *de novo* reconstruction of the transcriptome also revealed novel isoforms of RNAs that emerged in response to injury of the telencephalon. Moreover, our data suggest that the mRNAs of cholesterol synthesising and transporting proteins are subject to differential splicing thus contributing to the presumed adaptation of the cholesterol metabolism to the conditions in the injured brain (Figure 7B).

## Discussion

Unlike adult humans and other mammals, the adult zebrafish is able to efficiently repair injuries of the central nervous system. We analysed here the transcriptome for changes in the expression of mRNAs, their splice variants and regulatory RNAs including analysis of the targets of regulated miRNAs and transcription factors in response to injuries of the telencephalon. We noted profound changes in genes belonging to a large number of distinct cellular and physiological processes. As exemplified by the coordinated regulation of the cholesterol synthesizing enzymes and transporters, the genome responded in a multi-tiered manner with distinct and interwoven changes in expression of regulatory molecules to the physiological demands created by tissue damage and its repair. This multi-level regulation of the expression of cholesterol metabolising proteins uncovers an important process in the regenerating telencephalon. Our comprehensive analysis provides moreover an important source of information for future in-depth functional studies of specific genes and gene groups, regulatory molecules and splice variants in the regenerating zebrafish forebrain.

### Large scale response of the genome to telencephalon injury

The analysis of our sequencing data [14] with more than 600,000,000 reads from polyadenylated RNAs of control and injured telencephala revealed a change in expression of 4,989 genes. This represents 15% of all genes analysed and 29% of all genes detectably expressed in the samples. Thus, injury causes profound changes in the expression of information from the genome.

Brain regeneration is a complex process that entails many distinct physiological changes such as immune response, activation of glial cells, proliferation of stem cells, neurogenesis, axonogenesis etc. [20]. These previously described processes were all detected in our gene ontology analysis of protein coding genes adding an independent verification of our data and their analysis. In total, we scored 521 gene ontology terms and pathways with significant overrepresentation (adjp<0.05) in the transcriptome of the injured telencephalon relative to the uninjured control. These findings are in agreement with the large scale and complex demands of new proteins to cope with the inflicted injury. We had prepared cDNA from tissues at 5 dpl. At this time, the peak of proliferation of stem cells is reached [14]. Thus, the changes not only entail immediate early reactions to damage such as immune reaction but also genes with functions in repair of tissue function such as neurogenesis and axonogenesis.

Our findings on mRNA expression profiles are complementary to a recently published transcriptome study focusing on the immediate early changes in response to injury of the telencephalon [95]. Interestingly, this study reported activation of gene expression programs in both the injured and the uninjured hemispheres, even though the response was less pronounced and delayed in the injured hemisphere. In contrast, we never observed proliferation of stem cells or stem cell gene activation in the unlesioned hemisphere [10]. We systematically analysed hundreds of transcription regulators for their expression in the injured telencephalon by in situ hybridisation on sections. We did not observe gene activation in the uninjured hemisphere [14, 13, 95]. Demirci et al [95] inflicted lesions by inserting a needle into the nostril. In contrast, our protocol of injuring the telencephalon [96] involves the opening of the skull and inserting the needle directly into one hemisphere of the telencephalon. Most likely the protocol used by Demirci et al [95] causes damage of the second hemisphere or some of the extending nerves thereby causing activation of regenerative programs also in the seemingly uninjured hemisphere.

### Profound changes in splicing patterns in response to injury

The term “mRNA splice site selection” was also enriched among the genes with altered expression in the injured brain, - with 8 genes down-regulated in response to injury. This observation is in agreement with our systematic analysis of splice variants. We detected changes of splice patterns in 4,610 transcripts representing 1,309 genes. Thus, not only the overall levels of mRNAs were adapted to the physiological demands imposed by injury and repair but also the posttranscriptional processing of the mRNAs. In support, alternative splicing was reported for the modulation of the function of specific genes during neurogenesis in mammals [74, 59]. For example, in the developing mouse brain, the splicing factor PTBP2 targets mRNAs encoding DNM1 and modulates synaptic vesicle trafficking [64]. The mRNA isoforms were in most cases detected in both uninjured and injured telencephalic hemispheres. This suggests that injury causes a modulation of the function by shifting from one isoform to the other. Alternative splicing of mRNAs can also lead to the degradation of mRNAs [60]. Thus, alternatively, this shift of the predominant splice isoforms could thus be a means for adjusting the expression levels to the new physiological needs in the injured brain.

Taken together, our data suggest that alternative splicing represents another major response of the genome to cope with the physiological demands of the regenerating telencephalon. Since all splice variants were expressed in transcriptomes of controls and injured telencephala albeit at different levels, alternative splicing does not seem to control all-or-none effects but appears to be rather involved in the fine-tuning of the expression levels or functions of constitutively expressed genes.

### Alteration in cholesterol metabolism in response to brain injury

“Cholesterol biosynthesis” is a prominent gene ontology term among the genes whose expression was altered in response to injury. Cholesterol synthesis involves a pathway that initiates with the multistep synthesis of lanosterol from acetyl-CoA as precursor. Lanosterol is then converted by a whole battery of enzymes into 7-dehydrocholesterol and ultimately into cholesterol. With the exception of *fdps* mRNA, the mRNAs encoding cholesterol synthesizing enzymes of each of the steps from acetyl-CoA to cholesterol are down-regulated in the injured telencephalon. This suggests that cholesterol synthesis is co-ordinately reduced in response to injury.

Intriguingly, mRNAs encoding cholesterol transporters are elevated in the injured telencephalon. Transporters include also proteins associated with transport across endosomal membranes [49] suggesting alteration of cholesterol fluxes both across the plasma membrane and also within the cell into the endosomal compartment. Taken together, the regenerating telencephalon thus appears to systematically reprogram cholesterol metabolism from synthesis to relocation of cholesterol.

### Putative regulation of cholesterol synthesizing enzymes by Srebf2

In mammals, cholesterol synthesis is tightly regulated by posttranscriptional mechanisms involving the subcellular localisation of the Srebf2 transcription factor [25]. At high levels of available cholesterol, Srebf2is associated with Insig1 and Scap at the membranes of the endoplasmic reticulum (ER) and Golgi apparatus. Upon cholesterol shortage, this repressive association is dissolved and Srebf2 moves to the nucleus where it binds to the promoters of genes encoding the various enzymes of the cholesterol synthesis pathway and thereby induces the expression of the enzymes. In mammalian genomes, there are two related *Srebf* genes, *Srebf1* and *Srebf2*, - with Srebf2 being predominantly involved in regulation of genes encoding cholesterol synthesizing enzymes [24, 25, 26]. Similarly, the zebrafish genome harbours two *srebf* genes highly related to mammalian *srebf1* and *srebf2*.

Both Srebf1 and −2 are expressed in the adult zebrafish telencephalon. Our bioinformatic analysis of the 1-kb promoter upstream regions of genes encoding cholesterol synthesizing enzymes in the zebrafish genome revealed a strong enrichment of Srebf binding sites in the 1-kb promoter region of these genes. Also *insig1and scap* mRNAs are expressed in the zebrafish telencephalon. It is thus likely that the regulation of cholesterol synthesis involves relocation of Srebf2 protein from ER to nucleus in the zebrafish telencephalon similar to that seen in mammals.

Our comparative analysis of the injured and uninjured brain uncovered, however, in addition regulation of the *srebf2* mRNA level. *srebf2* mRNA was less abundant in the injured brain in agreement with the decreased expression of cholesterol synthesizing enzymes. This suggests an additional and distinct mode of regulation of Srebf2 activity besides the well-documented, posttranscriptional control of its activity through subcellular location as well documented in mammals. Intriguingly, we detected a Srebf2 binding site in the promoter of the *srebf2* gene suggesting auto-regulation via a positive feedback loop. Taken together, our *in silico* analysis suggests that the regulation of the level of Srebf2 mRNA is a potential mechanism how cholesterol synthesis is adjusted to the needs of the regenerating zebrafish telencephalon. In view of the expression of the Srebf2 co-factors *insig1* and *scap* in the telencephalon, it is, however, likely that Srebf2 activity is in addition regulated by posttranscriptional mechanisms and not just the abundance of the Srebf2 protein.

### microRNAs as additional regulatory mechanisms of cholesterol metabolism in the injured telencephalon

miRNAs are well known as regulators of large gene batteries [65]. A total of 184 miRNAs annotated in the zebrafish reference genome (GRCz11) were detectable by small RNAseq in the transcriptome of the adult zebrafish telencephalon. Of these, 31 miRNAs varied in level of expression upon injury. These miRNAs are distinct from miRNAs implicated previously in constitutive neurogenesis [61] and regeneration of the zebrafish optic nerve [27]. Thus, the scale and type of damage of the telencephalon may trigger specific responses both with respect to clearance of dead tissue, neurogenesis and regenerative processes. Given the fact that we prepared small RNAs from entire injured and uninjured hemispheres, it cannot be totally excluded that we failed to detect changes in miRNA expression in low-abundant cells such as stem cells and neuroblasts [1]. However, we detected constitutive expression of miR-9 which is expressed in neural stem cells in the telencephalon [79]. Our sensitivity of detection is thus very high and includes also stem-cell-specific miRNAs.

Potential targets of miRNAs were identified by the presence of the binding site of miRNAs in the 3’UTR of mRNAs expressed in the injured and uninjured telencephalon. Intriguingly, the expression of the miRNAs, *miR-182, miR-31, miR-155* and *miR-146a,* which were most strongly up-regulated in response to injury are all linked to the regulation of cholesterol synthesis. The *miR-182* seed sequence was found in the mRNAs encoding the enzymes Acat2, Hmgcs1 and Dhcrq24 of the cholesterol synthesis pathway. *miR-182* targets also the Srebf2 co-regulator *insig1* mRNA. Thus, *miR-182* affects cholesterol metabolism at two levels: i. The regulator Insig1 ii. Selected synthesizing enzymes. The seed sequences of*miR-31*, *miR-155* and *miR-146a* were present in the 3’UTR of five mRNAs coding for enzymes of the conversion of lanosterol into 7-dehydrocholesterol. In the mouse liver, depletion of *miR-155* results in an increase in hepatic level of cholesterol [62]. Similarly, *miR-146a* was shown to regulate the plasma level of cholesterol [63]. These observations in mice are consistent with the inferred role of these miRNAs in the injured zebrafish telencephalon.

In summary, these miRNAs could provide additional regulatory inputs that act in parallel with the Srebf2 factor on cholesterol synthesizing enzymes. Curiously, the three miRNAs, *miR-31, miR-155* and *miR-146a* target all one section of the cholesterol synthesis pathway, the conversion of lanosterol into 7- dehydrocholesterol (Figure 3A). This suggests that these may be key steps that need tight control.

### Alternative splicing as an additional mode of regulation of cholesterol metabolism

Alternative splicing that can increase the diversity of proteins [66] or leads to degradation of mRNAs or proteins [53] was noted for a number of genes involved both in synthesis and transport of cholesterol. In mammals, polypyrimidine tract binding protein 1 (PTBP1) splices mRNAs encoding several proteins of the cholesterol metabolism including the enzymes HMGCS1, FADS2, and PCSK9 [46, 47]. We found that the levels of mRNAs encoding the two zebrafish homologues Ptbp1a and Ptbp1b were significantly increased suggesting a role of Ptbp1a/b proteins in the regulation of cholesterol metabolism in the injured telencephalon of the zebrafish. In agreement, we observed a shift in the splice patterns of the zebrafish homologues of hmgcs1in response to injury. In mammals, this shift in splice patterns of hmgcs1 was paralleled by an overall decrease of the three proteins [47]. This suggests that the action of increased ptp1a/b leads to isoforms that are less stable, thereby contributing to the systemic decrease in the expression of mRNAs encoding cholesterol synthesizing enzymes in the injured telencephalon.

Another protein with alternatively spliced mRNA is Cadm1b that has a predicted Cholesterol 24-hydroxylase activity (GO term). Thus, in addition to decreased synthesis, increased degradation of cholesterol appears to be an activated process in the injured telencephalon.

Several mRNAs encoding transporters of cholesterol were also alternatively spliced after injury. These include mRNAs of the Low Density Lipoprotein Receptor *vldlr* which binds LDLs responsible for cholesterol transport through the blood stream [56]. Anxa6 together with NPC proteins mediates endosomal trafficking of cholesterol [49]. Two new donor sites which have not been annotated so far in the reference genome were discovered in exons of *anxa6* and *cadm1b* and were alternatively used in response to injury. Several transporter of the OSB family [57] were affected by alternative splicing. None of the alternatively spliced amino acid coding exons has a specific annotated function or structure in the InterPro database. It remains thus to be seen whether these alternatively spliced proteins have altered properties such as function, stability or subcellular locations. Taken together our results suggest that alternative splicing is an important process contributing to the altered expression/activity of cholesterol metabolising proteins in response to injury.

### Relevance of modulation of cholesterol metabolism

Key questions are why cholesterol metabolism is so tightly regulated and why the down-regulation of it is a feature of the injured telencephalon.

Cholesterol is an essential component of many cellular processes. It determines the biophysical and biochemical properties of membranes. It is a precursor of steroid hormones, and cholesterol derivatives are important secondary modifications of proteins such as Wnt receptors [67] or Hedgehog ligands [68]. Thus and given its general hydrophobic property, free excess cholesterol has an impact on the functional integrity of membranes and the communication between cells. Debris from damaged axonal processes, in particular their membrane-rich myelin sheaths, may likely be abundant sources of extracellular cholesterol. It is probably very critical for cell survival and efficient repair of the brain to control free cholesterol levels very tightly.

The coordinated down-regulation of expression of genes encoding cholesterol synthesizing enzymes and the up-regulation of transporters suggests that the injured telencephalon switches from synthesis of cholesterol to its import from the extracellular environment. In addition, the up-regulation of genes coding for cholesterol transporters (Npc1, Npc2), which are linked to endosomes [49] may reflect an additional switch to cholesterol storage or degradation rather than synthesis. In this context, it is important to note that the blood brain barrier prevents efficient exchange of cholesterol between the brain and the rest of the body [50], possibly necessitating this tight control within the brain. In the injured brain of mice [75], cholesterol 25-hydroxylase levels are increased in microglial cells. This enzyme converts cholesterol into a more hydrophilic and allows there by its crossing of the blood brain barrier and degradation in the liver. We observed an elevated level of cholesterol 25- hydroxylase mRNA in the injured zebrafish brain suggesting that increased efflux to the liver may also be a mechanism to reduce cholesterol levels in the injured zebrafish brain. Furthermore, in the injured mouse and rat brain, APOE, a transporter of cholesterol was increased upon injury [69, 70].

Thus, also in the mouse, cholesterol degradation and transport appears to be increased upon injury. To our knowledge, however, the systemic regulation of cholesterol metabolizing enzymes which we observed in the injured zebrafish telencephalon was not reported so far for the mammalian brain. This control of cholesterol metabolism may be of medical relevance. It may open possibilities to combat the outcomes of conditions like stroke, injury and neurodegeneration in the human brain. Interestingly, decreasing the level of circulating cholesterol in the rat improves the recovery after brain injury [70]. Taken together, our comparative *in silico* analysis of the transcriptomes of the injured and uninjured telencephalon of the adult zebrafish suggests that regulation of cholesterol levels is an important process for brain regeneration.

### Multi-layer regulation of cholesterol metabolism a means of robustness or a mood of evolution?

Cholesterol metabolism is regulated at multiple levels in response to injury. At the level of inferred protein functions, an overall switch from synthesis to transport and possibly also storage and degradation of cholesterol is evident from the comparative analysis of the transcriptomes of the injured and uninjured telencephalon. When the putative regulatory mechanisms were explored, changes of expression of regulatory molecules suggested multiple synergistic and complementary regulatory networks controlling cholesterol synthesis and transport. These include a decrease of Srebf2 mRNA leading to reduction of the key transcription activator of most enzymes of the cholesterol synthesis pathway. Drawing from the mammalian literature [42], the expression of key regulators (Insig1, Scap) in the zebrafish telencephalon and given the general high conservation of many regulatory mechanisms between fish and mammals, it is likely that the activity of zebrafish Srebf2 is also regulated by subcellular distribution. Thus, in addition to down-regulation of the *srebf2* mRNA, the corresponding protein is likely not located in the nucleus in the injured telencephalon. Changes in splice patterns of cholesterol synthesizing enzymes and transporters may alter protein function or lead to degradation of the mRNA or encoded enzymes adding another principle of regulation. Further layers of regulation are conferred by changes in expression of regulatory RNAs. Up-regulation of miRNAs targeting the mRNAs of a subgroup of cholesterol synthesising enzymes contribute to the decrease of the target RNAs. Changed expression of lncRNAs overlapping with several genes encoding cholesterol synthesising or transporting proteins offer yet other layers of regulatory principle woven into the control of cholesterol metabolism.

A key question is why cholesterol metabolism requires such a complex multi-layered control. The transcriptional changes in cholesterol metabolizing genes and their multilevel regulation may be a reflection of the brain’s autonomy with respect to cholesterol metabolism. The crucial functions of cholesterol and the pathogenic effects of excessively high cholesterol levels may call for efficient and robust mechanisms. This robustness may be best achieved by complementary and synergistic modes of regulation. Alternatively, this architecture of regulatory mechanisms may be a reflection of how living systems evolve. By randomly recruiting and adapting components of the cells existing repertoire of gene regulatory mechanisms, this seemingly rather complex regulatory network architecture may have arisen. As the evolved mechanisms were effective, they were maintained. Thus, this complexity most likely reflects both evolutionary process and robustness in adaptation of cholesterol levels to the physiological state during injury and repair of the brain.

## Conclusion

The transcriptome of the adult zebrafish telencephalon reacts to injury with profound changes in mRNA expression including shifts in mRNA splice patterns and expression of regulatory RNAs (miRNAs, lncRNAs). These large scale changes of retrieval of nuclear information affect many cellular processes and pathways. As an example, our data support repression of cholesterol synthesis and a redistribution of existing pools of cholesterol in the injured telencephalon. Multiple and distinct modes of gene regulation appear to converge on the cholesterol pathway presumably to cope effectively with excess cholesterol in the injured telencephalon. Thus, regulation of cholesterol appears to be a key event of the regenerating zebrafish telencephalon. These observations may also have medical relevance.

## Material and methods

### RNASeq data analysis

RNASeq data were generated as described previously [14]. Reads were mapped against the zebrafish reference genome GRCz11 with STAR [31]. For reads mapped at multiple loci, only the mapping with the highest quality score was outputted. The purpose was to more accurately quantify the expression of genes. Raw read counts at gene level were computed with HTSeq in union mode [32]. Expression normalization and differential expression analysis were both carried out with DESEq2 [33]. Aberrant values of expression were flagged and corrected with a generalized linear model with DESeq2. Taken into account the high depth of sequencing, i.e. greater than 200,000,000 on average per sample, a gene was considered as expressed with an average normalized level of expression across all samples greater than 100. Differences in expression between control and injured telencephalic hemispheres were assessed also with DESeq2. The p-values of the Wald tests were adjusted with the Bonferroni method. A threshold of 0.05 was applied on adjp to identify significant changes in expression in response to injury. No threshold was applied on FC to exhaustively identify differentially expressed genes.

lncRNA genes transcribed in the adult zebrafish telencephalon were identified based on the tag “biotype” extracted from the annotation of the zebrafish reference genome. The annotation of the neighboring genes directly upstream and downstream, with no threshold of distance, was carried out with the R packages GenomicRanges [34].

For the functional annotation of the zebrafish genome, the latest gene ontology terms [21], signaling pathways [22] and metabolism pathways [23] were retrieved from their respective database. The enrichment was then tested with the one-tailed exact Fisher test, as previously published [35]. The Fisher p-values were corrected with the False Discovery Rate (FDR) method as previously published [35]. A threshold of 0.05 was applied on corrected p-values to identify significantly enriched biological functions.

The two binding motif of SREBF2 identified in vertebrates were retrieved from the database JASPAR 2018 [36] and were mapped, with HOMER [37], in the promoter sequence of all genes with significant variation in the level of transcripts in response to injury. The promoter region was defined as 1-kb upstream of the transcription start site, provided by the annotation of the zebrafish reference genome. For the mapping of SRE with HOMER, a background set was created with the same number of sequence, i.e. 4,989, randomly extracted from the zebrafish reference genome GRCz11. The background sequences were also of the same size, i.e. 1 kb. Both forward and reverse strands were analyzed.

To investigate alternative splicing of polyadenylated RNAs, transcripts synthesized in the adult zebrafish telencephalon were first *de novo* reconstructed from mapped RNASeq with STAR. The mapping of the reads at splicing junction was refined with a second pass taken into account splicing junctions identified in both control and injured RNASeq samples. From the mapped reads, transcripts were *de novo* reconstructed, with Leafcutter [39], with no limits in the number of introns per transcript and novel splice junctions supported by a minimum of 20 split reads. For each transcript differential splicing between control and injured telencephalic hemisphere was assessed with Leafcutter as well. The p-values were corrected with the False Discovery Rate method as recommended. Significant alternative splicing of transcripts in response to injury were identified with two parameters: 1. applying a threshold of 0.05 on adjusted p-values, 2. the corresponding splicing junction was covered by at least 20 mapped reads. Results were visualized with the genome browser IGV [40] and transcript isoforms were manually reconstructed. The sequence of spliced exons were retrieved from Ensembl [80] and the corresponding protein domains were identified with the software InterPro [55] relying on annotation of protein domains present in the database UniProt [76].

### Sequencing of small RNAs and microRNA analysis

Small RNA libraries were prepared from 1 μg of total RNAs with the Small RNA Library Preparation kit (Illumina) following the manufacturer’s protocol. Three libraries for control and injured telencephalic hemispheres were sequenced with a HiSeq1500 (Illumina). The adaptor sequence (Illumina) was trimmed from raw reads with Cutadapt [99] for a final insert size of 21, 22 or 23 nucleotides.

Passing all quality controls carried out with FASTX toolkit (http://hannonlab.cshl.edu/fastx_toolkit/index.html), reads were mapped against the zebrafish reference genome GRCz11 with STAR [31]. No soft-clippings were allowed, only one mismatch was allowed and only mappings with a quality of 30 (Phred score) were outputted. Raw read counts were computed with HTSeq [32] in union mode and with an annotation file including all known miRNA loci in the annotation of the zebrafish reference genome GRCz11, as recommended by the ENCODE project [41]. Expression normalization and differential expression analysis were both carried out with DESEq2 [33], as described above. A threshold of 0.05 was applied on adjusted p-value to identify significant changes in steady state levels of miRNAs upon injury. To identify strong changes in levels of miRNAs upon injury, thresholds of 0.25 and 2 were applied on FC. A miRNA was considered as expressed with an average normalized level of expression across all samples greater than 10. Predicted target mRNAs, specific for the zebrafish, were retrieved from the data base TargetScanFish [42]. No filters were applied on the tissue where the miRNAs were originally expressed.

### Preparation of biological samples and qRT-PCR

Injury was inflicted to the telencephalon as described previously [1]. For qRT-PCR, total RNA was isolated from injured and uninjured telencephalic hemispheres using Trizol (Life Technology). First strand cDNA was synthesized from 1 μg of total RNA with the Maxima First Strand cDNA kit (Thermo Scientific) and according to the manufacturer’s protocol. qRT-PCR was carried out with a StepOnePlus Real-time qRT-PCR system (Applied Biosystems) and SYBR Green I fluorescent dye (Promega). Expression levels of genes were normalized to β- actin expression and the relative expression levels were calculated using the 2^-ΔΔCT^ method. Real-time qRT-PCR was carried out in triplicates of independently prepared samples and repeated once. Differences in relative expression between control and injured telencephalic hemispheres were tested with the one-tailed t-test.

## Declarations

Availability of data and materials: mRNAseq and small RNAseq data are available on the Gene Expression Omnibus data base under the accession identifiers GSE161137 and GSE160992, respectively.

## Competing interests

The authors declare no competing interests.

## Funding

We are grateful for support by Helmholtz Association BioInterfaces_TM Program,

## Authors’ contributions

US, OA and SR designed the study. OA carried out the sequencing experiments. LL and ND tested the reproducibility of the results. VG analyzed and integrated the results. US, VG and OA interpreted the results. US and VG wrote the manuscript. All the authors read and approved the manuscript.

## Acknowledgements

We thank Masanari Takamiya for his valuable comments, Tanja Both for preparing the sequencing libraries and Martin März for injuring telencephala.

## Abbreviations

adjp: adjusted p-value
cDNA: complementary deoxyribonucleic acid
ChIP: chromatin immune-precipitation
CNS: central nervous system
dpl: day post lesion
ER: endoplasmic reticulum
FC: fold change
FDR: false discovery rate
GO: gene ontology
hpl: hour post lesion
kb: kilobase
KEGG: Kyoto encyclopedia of genes and genomes
LDL: low density lipoprotein
mRNA: messenger RNA
miRNA: microRNA
lncRNA: long non-coding RNA
NSC: neuronal stem cell
OSB: oxy sterol binding
qRT-PCR: quantitative reverse transcription polymerase chain reaction
RGC: radial glial cell
RNA: ribonucleic acid
SRE: sterol regulatory element
SREBF: sterol regulatory element binding family
TF: transcription factors
UTR: untranslated region

**Supplementary Figure: Reconstruction of alternatively spliced isoforms of transcripts related to cholesterol metabolism.** Solid square: annotated exon; dashed square: novel exon; red: increased usage of junction; green: decreased usage of junction; 5’: 5’UTR; *: stop codon; number: Ensembl exon identifier.

**Supplementary Table 1: Polyadenylated RNAs with significantly changed levels upon injury.**

**Supplementary Table 2: Significantly enriched biological functions among differentially expressed genes.**

**Supplementary Table 3: Differentially expressed genes encoding proteins with function related to cholesterol metabolism.**

**Supplementary Table 4: Comparison of genes with cholesterol biosynthetic function in the human and zebrafish genome and with mapped SRE motifs in the 1-kb promoter.**

**Supplementary Table 5: SRE motifs mapped in the promoter of genes with functions related to cholesterol biosynthesis in the zebrafish genome.**

**Supplementary Table 6: Genes differentially expressed in response to injury with SRE motif(s) in the 1-kb promoter region.**

**Supplementary Table 7: microRNAs with significant changes in level upon injury.**

**Supplementary Table 8: Significant alternative splicing in response to injury.**

**Supplementary Table 9: Significantly enriched biological functions among differentially expressed genes harboring the SRE motif in their promoters.**

## Notes

### Competing Interest Statement

The authors have declared no competing interest.

